# Non-invasive investigation of the morphology and optical properties of the upside-down jellyfish *Cassiopea* with optical coherence tomography

**DOI:** 10.1101/2023.01.10.523435

**Authors:** Niclas Heidelberg Lyndby, Swathi Murthy, Sandrine Bessette, Sofie Lindegaard Jakobsen, Anders Meibom, Michael Kühl

## Abstract

The jellyfish *Cassiopea* largely cover their organic carbon demand via photosynthates produced by their microalgal endosymbionts, but how holobiont morphology and optical properties affect the light microclimate and symbiont photosynthesis in *Cassiopea* remain unexplored. Here, we use optical coherence tomography (OCT) to study the morphology of live *Cassiopea* medusae at high spatial resolution. We include detailed 3D reconstructions of external micromorphology, and show the spatial distribution of endosymbionts clustered in amoebocytes and white granules in the bell tissue. Furthermore, we use OCT data to extract inherent optical properties from light scattering white granules in *Cassiopea* and show that white granules enhance local light availability for symbionts in close proximity. Individual granules had a scattering coefficient of μ_s_ = 200-300 cm^-1^, and a scattering anisotropy factor of *g* = 0.7, while large tissue regions filled with white granules had a lower μ_s_ = 40-100 cm^-1^, and *g* = 0.8-0.9. We combined OCT information with an isotopic labelling experiment to investigate the effect of enhanced light availability in whitish tissue regions. Algal symbionts located in whitish tissue exhibited significantly higher carbon fixation as compared to symbionts in anastomosing tissue (i.e., tissue without light scattering white granules). Our findings support previous suggestions that white granules in *Cassiopea* play an important role in the host modulation of the light-microenvironment.

## Introduction

Characterization of anatomical structures and morphology in photosymbiotic animals is important to understand the interplay between host light-modulation and the spatial organization of photosymbionts in complex holobionts, such as corals and other photosymbiotic cnidarians. External structures can be relatively easily identified using macro- and microscopic imaging in the visible wavelength range on intact individuals with various levels of preparation (Shapiro et al., 2016; Sivaguru et al., 2014). However, imaging deeper into tissue is challenging with conventional or laser scanning microscopes, where the observational depth is typically limited to a few hundred micrometers before absorption and scattering of light reduces image quality substantially, and fluorescent labelling of structures is often necessary to achieve sufficient contrast (Salih, 2012). Invasive tissue clearing of fixed specimens can enhance light penetration for 3D reconstruction of intact whole organisms (Dekkers et al., 2019). However, serial sectioning in combination with light or electron microscopy is often required to obtain a combination of macrostructural observations and detailed information of internal organization (Colley and Trench, 1985). This is both destructive and slow, and involves tissue sample preparation steps that can create artefacts in tissue organization (Huebinger et al., 2018; Loussert-Fonta et al., 2020).

In this context, optical coherence tomography (OCT) presents an attractive alternative imaging modality (Fercher et al., 2003; Huang et al., 1991; Schmitt, 1999), as it enables non-invasive *in vivo* imaging of live specimens using near infrared (NIR) wavelengths, allows for high-resolution tomographic scanning of tissue in real time, and enables 3D reconstruction of large samples with a potential depth of penetration of several millimeters depending on the scattering properties of the sample (Schmitt, 1999; Wangpraseurt et al., 2017b). Previous studies have utilized OCT for visualizing and quantifying structural characteristics, such as surface area, volume, and porosity in biofilms (Depetris et al., 2021; Wagner et al., 2010), terrestrial plants (Hettinger et al., 2000), and marine invertebrates (Speiser et al., 2016), including scleractinian corals (Jaffe et al., 2022; Wangpraseurt et al., 2017b; Wangpraseurt et al., 2019).

The OCT measuring principle is analogous to ultrasound imaging and is based on the detection of directly backscattered light from refractive index mismatches in a sample, measured as the echo time delay and intensity of directly backscattered light from internal microstructures in materials or tissues (Fujimoto et al., 2000). OCT imaging uses low coherent NIR light to generate tomographic data composed of axial point measurements (A-scan) that are used to generate two-dimensional cross-sectional images (B-scan) or three-dimensional data cubes (C-scan) of backscattered photons using interferometry. Besides providing structural information from tomographic measurements of relative backscatter intensity, OCT systems can also be calibrated for absolute measurements of reflectivity. In combination with a theoretical bio-optical model based on an inverse Monte Carlo method (Levitz et al., 2010), calibrated OCT datasets can be used to extract optical properties, i.e., the scattering coefficient, μ_s_ (mm^-1^), and anisotropy of scattering, *g* (where *g* = 1 indicates completely forward scattered, and *g* = 0 indicates isotropic scattering). More details on the basic principle of OCT can be found elsewhere (Fercher et al., 2003; Huang et al., 1991; Schmitt, 1999). Furthermore, we note that OCT can also be combined with spectroscopic measurements to obtain a wider range of optical parameters and higher spatial resolution (Spicer et al. 2019).

The optical properties in corals have received increased attention over the last decade, and both coral tissue and skeleton have been found to exhibit strong light scattering properties (Enríquez et al., 2005; Enríquez et al., 2017; Lyndby et al., 2016; Spicer et al., 2019; Wangpraseurt et al., 2012; Wangpraseurt et al., 2014). Host-produced fluorescent pigments (FPs) in particular enable cnidarians to enhance scattering of incident light in the tissue, and the backscatter of light from the coral calcium carbonate skeleton further enhances light capture and photosynthesis by the endosymbiotic dinoflagellate algae (Enríquez et al., 2005; Lyndby et al., 2016; Marcelino et al., 2013; Swain et al., 2016; Terán et al., 2010; Wangpraseurt et al., 2014; Wangpraseurt et al., 2017a). Wangpraseurt et al. (2017b) used OCT imaging for real-time quantification of tissue movement in coral polyps, providing insights into symbiont and FP movement and density. They demonstrated how optical properties not only differ between tissue regions, but that tissue movement can lead to dynamic modifications of local optical properties of the coral tissue, potentially enhancing or reducing photon absorption (Wangpraseurt et al., 2019). However, we are not aware of such use of OCT for investigating other cnidarians.

The so-called up-side-down jellyfish *Cassiopea* is currently promoted as an important model system for cnidarian photo-symbiosis (Medina et al., 2021; Ohdera et al., 2018). *Cassiopea* hosts photosynthetic algae belonging to the same dinoflagellate family (Symbiodiniaceae) as symbionts in scleractinian corals (Lampert, 2016). Adult *Cassiopea* medusa live mostly in shallow benthic environments, where they position themselves up-side down on the sediment basking in the sun to expose their photosynthesizing algae that supplement the carbon demand of the heterotrophic host with autotrophically acquired carbon (Freeman et al., 2016; Lyndby et al., 2020; Welsh et al., 2009). The symbiont algae are found concentrated in the host tissue on the oral side of the bell and in oral arms of the medusa (Colley and Trench, 1985; Estes et al., 2003). *Cassiopea* exhibit a range of colors from purple-blue to reddish (Lampert, 2016; Lampert et al., 2012) but most commonly have a blue pigmentation associated with the bell disc and oral vesicles (Blanquet and Phelan, 1987), which was recently identified as a new group of pigments, i.e., Rhizostomins (Lawley et al., 2021). Another common trait of adult *Cassiopea* is the whitish appearance of certain tissue regions containing granules, which is most pronounced in the oral arms and stretches radially from manubrium to the bell margin and rhopalia (Bigelow, 1900). The function of the white granules in *Cassiopea* is still unknown, but it has been suggested to play a role in the mitigation of photodamage and enhancement of photosynthesis (Blanquet and Phelan, 1987; Phelan et al., 2006), where the white granules and colored pigments in *Cassiopea* might serve similar functions to that of the skeleton and host pigments in corals.

Here, we used non-invasive OCT imaging to visualize the structure of live *Cassiopea* sp. medusae and provide details of specific anatomical features in intact, living specimens as well as cut-out tissue samples from living specimens. We identified two highly heterogeneous tissue regions in the bell, namely the rhopalia canals containing a high density of white granules and the anastomosing tissue without white granules visibly present. Optical properties were extracted from these regions using the OCT data, and a labelling experiment using stable isotopes of ^13^C-bicarbonate and ^15^N-ammonium was conducted to test if the different optical properties altered photosynthetic performance of algal symbionts found in the two tissue regions.

## Methods

### *Cassiopea* husbandry and preparation

*Cassiopea* sp. medusae acquired from DeJong Marinelife (Netherlands) were cultivated at the Marine Biology Section (MBS), University of Copenhagen (Helsingør, Denmark) in a 50 L aquarium. Medusae were fed 5 times per week with recently hatched *Artemia* nauplii and supplemented with AF Amino Mix (Aquaforest) 2-3 times per week. Medusae were kept at 25°C in artificial seawater (ASW) with a salinity of 35 and a pH of 8.2. Water was recycled inside the tank using a small internal filter, as well as UV light filtration. Illumination was maintained with a programmable LED aquarium lamp (Triton R2, Pacific Sun) running a 12/12-hour day/night cycle with a downwelling photon irradiance (400-700 nm) of 350 μmol photons m^-2^ s^-1^, as measured just above the water surface using a calibrated spectroradiometer (MSC-15, GigaHertz-Optik). We used *Cassiopea* medusae in a size range from 7 mm up to several cm in diameter to scan the internal and external anatomy of the *Cassiopea* medusa life stage.

### Optical Coherence Tomography

We used a spectral domain OCT imaging system (Ganymede II, Thorlabs GmbH) comprised of a base unit with a non-coherent NIR light source (930nm; GAN611) and a scanning system (OCTG9) fitted with either the OCT-LK3-BB lens kit (field of view 10×10 mm, lateral resolution 8 μm, effective focal length 36 mm) for complete medusa scans, or an OCT-LK2-BB kit (field of view 6×6 mm, lateral resolution 4 μm, effective focal length 18 mm) for closeup scans with a higher resolution. The axial resolution was 5.8 μm. For imaging, animals and cut-out tissue samples were kept in a small petri dish submerged in either filtered ASW or 100% glycerol, typically with 4 mm liquid on top of the tissue surface. Prior to the OCT imaging, animals were anaesthetized with MgCl_2_ (50% w/v in ASW) that was added dropwise to the petri dish until the individual stopped pulsating. Cut-out tissue, if not stable in a favorable orientation relative to the OCT system, was fixed with hypodermic needles onto a cork surface covered with dark-cloth to reduce light reflection.

### Extraction of optical parameters using OCT

Inherent optical properties like the scattering coefficient, μ_s_ [cm^-1^], and the anisotropy of scattering, *g*, can be estimated from OCT scans of biological tissues using theoretical models of light propagation based on the inverse Monte Carlo method (Levitz et al., 2004; Thrane et al., 2005). A more detailed description of optical parameter extraction from OCT scans of tissue can be found elsewhere (Fercher et al., 2003; Wangpraseurt et al., 2019).

Briefly, OCT B-scans were acquired with a resolution of 581×1024 pixels, over a fixed depth interval of 2.8 mm, and variable distance in the X-plane. The setup was optimized to yield the highest signal at a fixed distance of 0.4 mm from the top of the scan. The OCT reflectivity (R) was calibrated before measurements using homemade reflectance standards with an immersion oil-glass, a water-glass, and an air-glass interface. R values from the standards were determined using Fresnel’s equation:

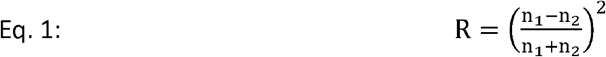

using the refractive index (n) for air (1), water (1.33), immersion oil (1.46), and quartz glass (1.52). The OCT signal (in decibel, dB) from the samples, was then converted to the depth-dependent R via a linear fit of log_10_(R) versus OCT intensity values (see Wangpraseurt et al., 2019 for details).

Calibration of the focus function of the objective lens was performed by measuring the OCT signal fall off, in steps of 0.1 mm, from either side of the focal plane (z = 0.4 mm) to z = 0 and 0.8, respectively. The signal loss from the focal plane follows an exponential decay function. The determined R values from the sample scans were corrected by dividing with the exponential fit. Then the corrected R values were plotted against sample depth (z, distance from focal volume) and fitted to the exponential decay function:

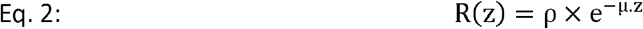

where ρ (dimensionless) is the light intensity and μ is the signal attenuation (cm^-1^) from the focal volume. The fit was considered satisfactory if R^2^ > 0.5. The first few pixels from each scan were excluded due to high reflectivity at interface, arising from the refractive index mismatch between the water and epidermis.

Using the grid method (Levitz et al., 2010), values of ρ and μ were matched to *g* and μ_s_ based on the theory described in previous studies (Samatham et al., 2008; Wangpraseurt et al., 2019). It was assumed that the *Cassiopea* tissue absorption at 930 nm was negligible, and the absorption was dominated by water (μ_a_ ∼ 0.43 cm^-1^). The effective numerical aperture (NA) was 0.11. The refractive index of the investigated tissue was estimated to be 1.473 (see results and discussion).

The targeted areas for extraction of optical properties in *Cassiopea* tissue were i) bell tissue with white granules (along rhopalia canals), and ii) symbiont clusters in anastomosing tissue. Scans in ASW were done on intact tissue, and on cut-out tissue where the epidermis was removed by gently peeling the epidermis from the underlying bell using tweezers.

Raw OCT files, and selected figures presented as 3D animations (figures 1-6) can be found in the supplementary information.

**Figure 1.**
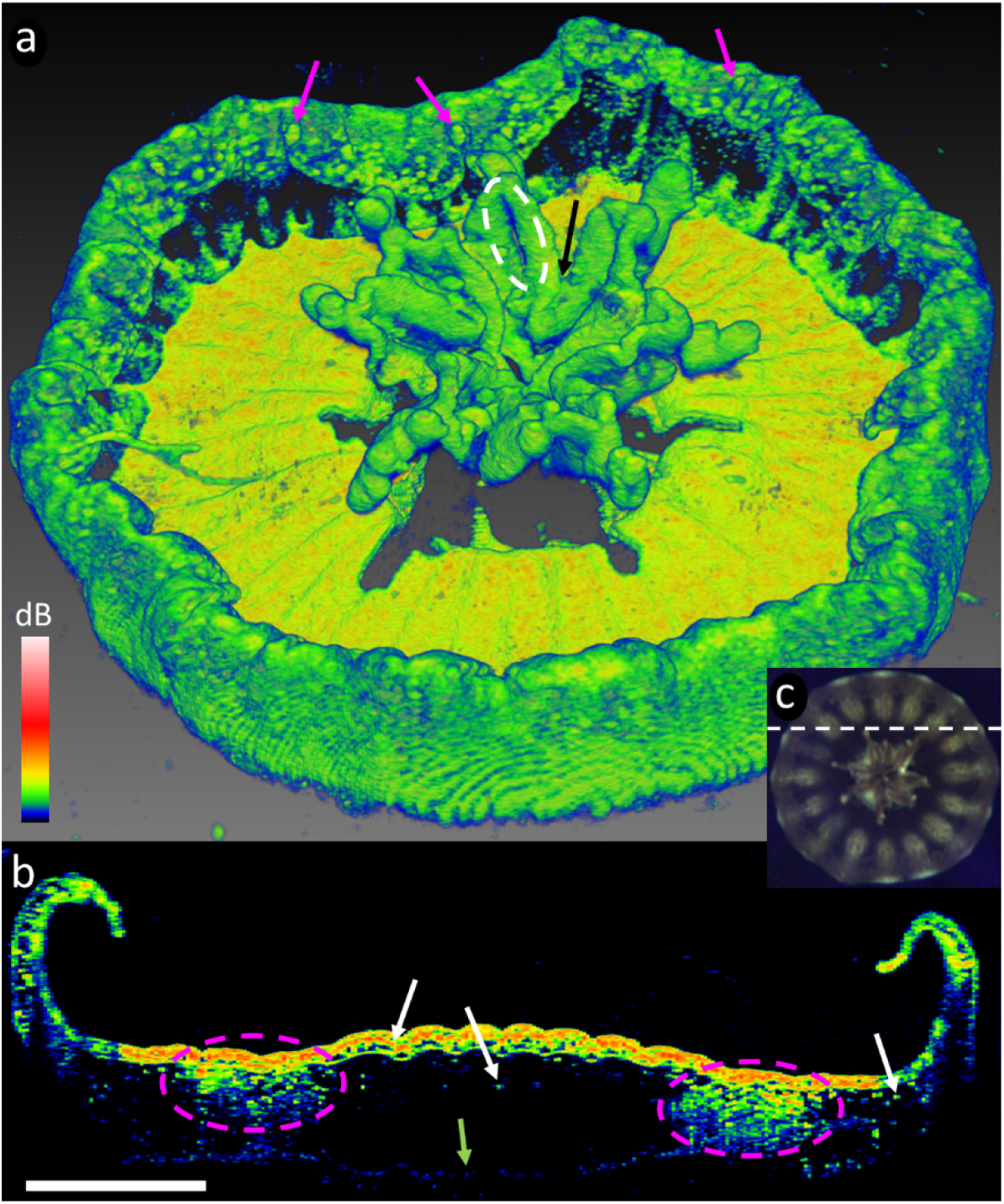
(a) 3D scan (C-scan) of an entire juvenile *Cassiopea* sp. medusa, using a FOV of 7×7×2.8 mm (X-Y-Z), a resolution of 350×350×1024 pixels, corresponding to a voxel size of 20×20×2.74 μm. Rhopalia can be seen along the bell rim (purple arrows), and first oral arm bifurcation (black arrow) and an oral grove (white dashed circle) are clearly visible on the oral arms. (b) Tomographic cross section (B-scan) in the XZ plane through the bell, revealing the *in vivo* distribution of dense symbiont clusters underneath the subumbrella epidermis and diffusely spread deeper in the bell (white arrows), and large patches of white granules (purple dashed circles) underneath epidermis. A faint line of the exumbrella epidermis is visible at the bottom (green arrow). (c) Overview photo of the animal under OCT observation, the dashed white line indicates the B-scan presented in (b). White scale bar represents 1 mm. Colored scale bar indicates relative OCT signal strength. An animated 3D representation (Movie 1) and the raw OCT file can be found in the supplementary information.

**Figure 2.**
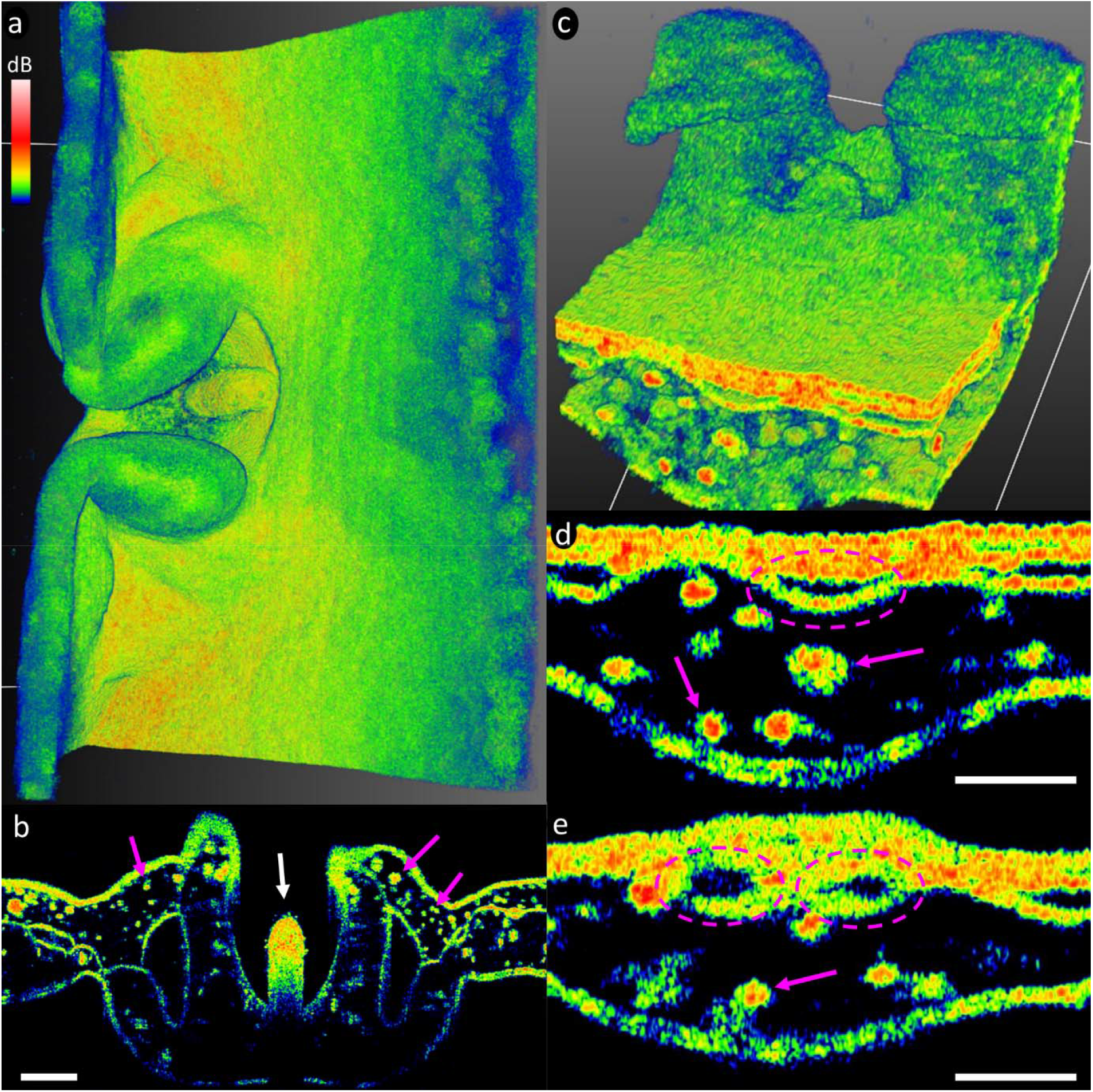
Rhopalia of *Cassiopea* medusae of ∼4 cm (a) and ∼1 cm (c) in diameter. (b) The rhopalium is highly scattering of NIR light and appears almost solid (white arrow). (d) The rhopalia canal can be observed running out to the base of the rhopalia (purple dashed circles), before splitting into two canals that reach out into the marginal lappets on each side of the rhopalia (e). Throughout the scans, clusters of symbionts and white granules were diffusely spread out in the bell margin and marginal lappets along the rhopalia canal and into the bell margin (purple arrows). Scale bars represent 200 μm in panel b, and 100 μm in d-e. Colored scale bar indicates relative OCT signal strength.

**Figure 3.**
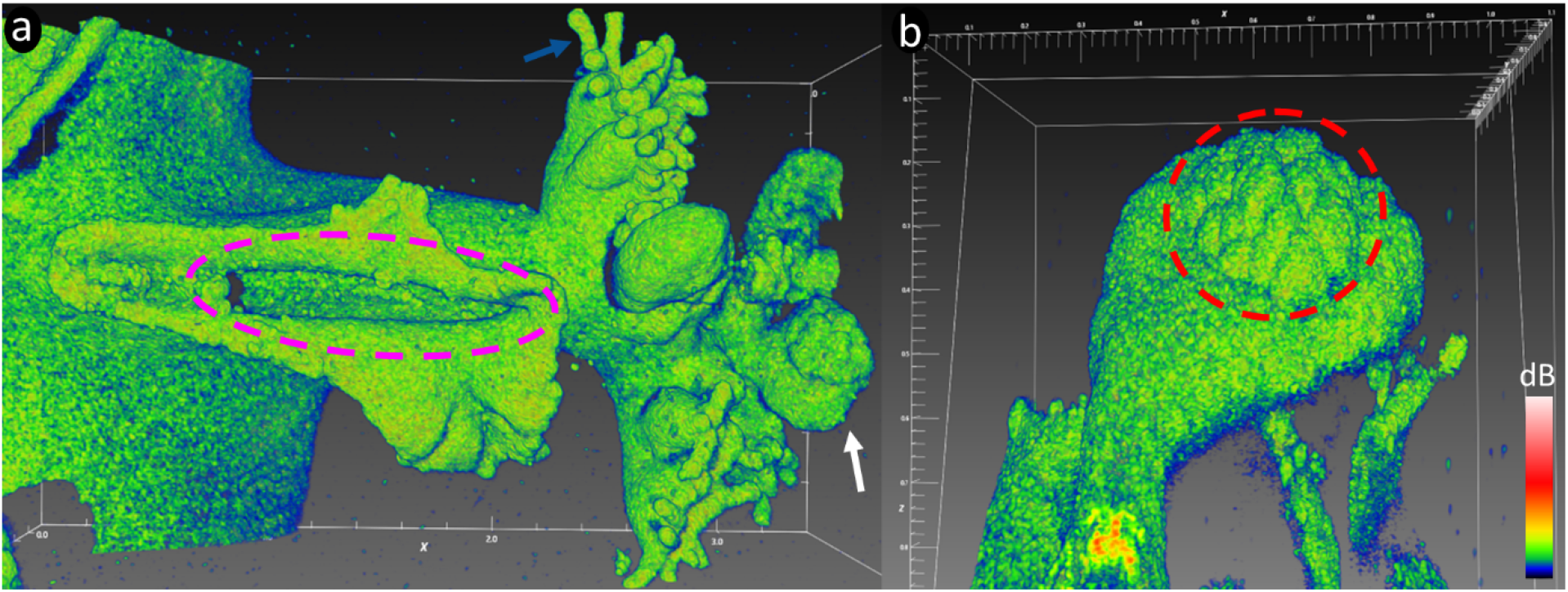
(a) Oral arm with oral groove (purple dashed circle; ∼1 mm in length). Sectional scans of the arm showed that the oral groove extended into the arm and further into the manubrium (not shown here). Oral vesicles (white arrow) were found apically on the arm, along with secondary mouths covered in frigid digitata (blue arrow). (b) closeup of vesicle with a developing cluster of cassiosomes (red dashed circle). White scale bars in mm. Colored scale bar indicates relative OCT signal strength.

**Figure 4.**
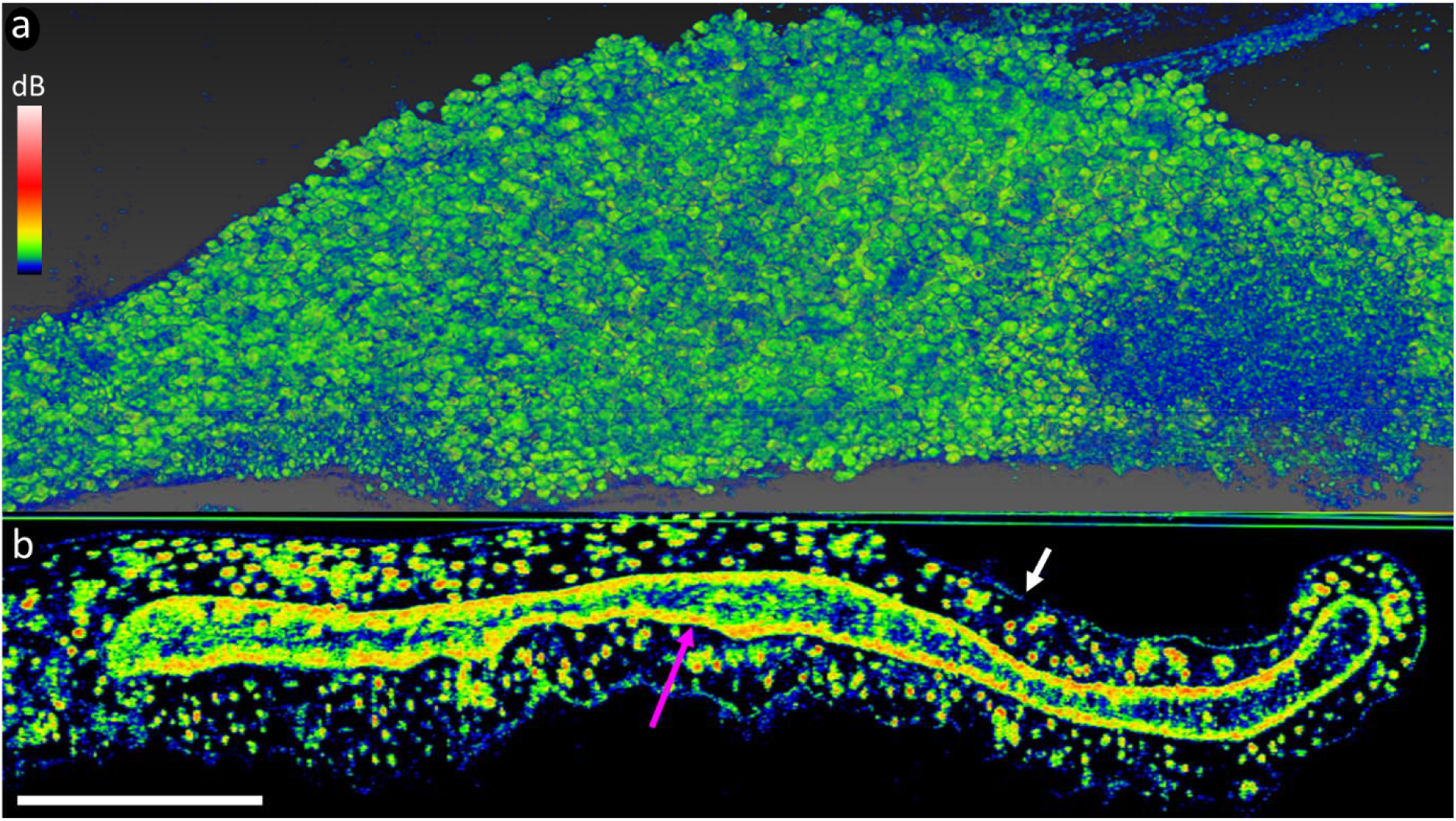
(a) Overview of an intact vesicular lappet submerged in pure glycerol. The refractive index of glycerol and *Cassiopea* epidermis match closely, making the epidermis near-invisible and revealing underlying dispersed clusters of symbionts in the mesoglea (spherical shapes dotted throughout the sample). (b) Cross-sectional scan of the lappet. The epidermis was still faintly visible (white arrow). The endodermis (purple arrow) was much more distinct compared with images in water, due to the refractive mismatch between the endodermis and mesoglea in the lappet. The lappet clearly exhibits a closed cavity that does not extend back into the oral arm. What this cavity consists of, or if symbionts are present inside, remains to be studied in further detail. White scale bar represents 500 μm. Colored scale bar indicates relative OCT signal strength.

**Figure 5.**
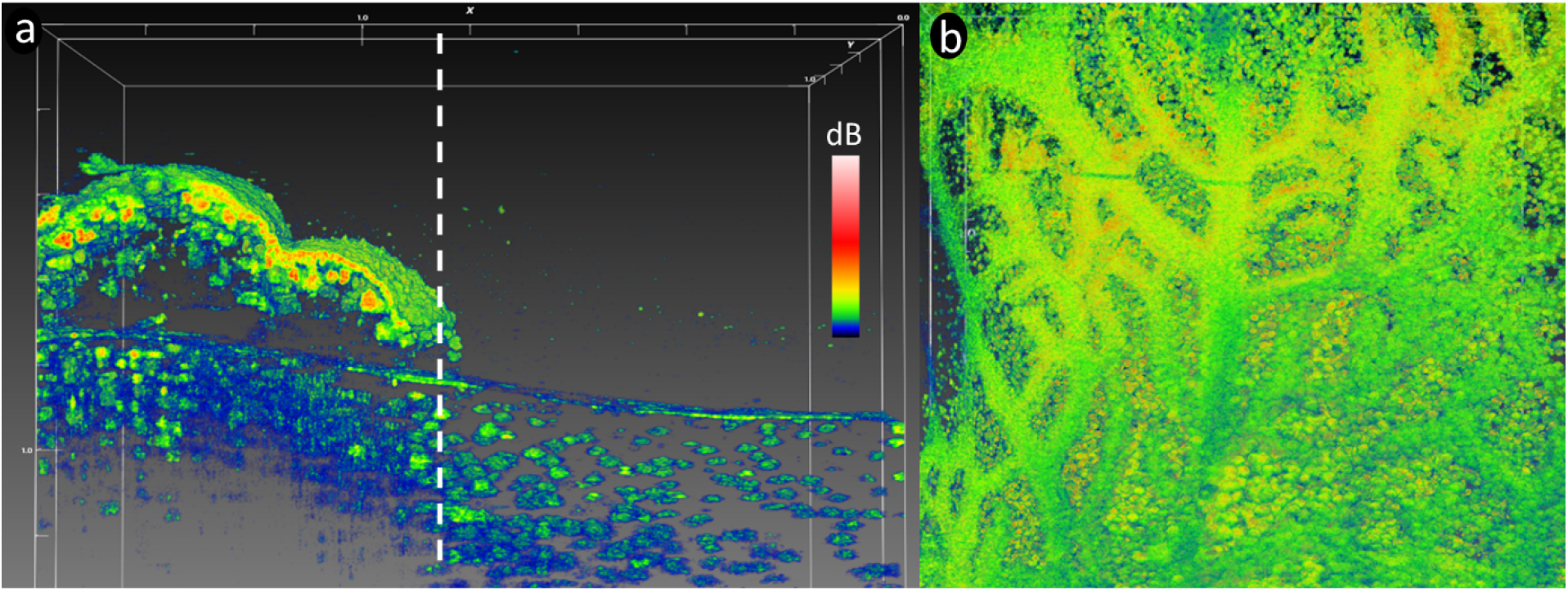
(a) OCT scan of bell tissue with part of the subumbrella epidermis and mesoglea removed (right side of dashed line), improving scan penetration depth and clarity in this area. (b) Overview of bell tissue with subumbrella layer removed. The remaining gastrodermis from the radial canals was still attached to the exumbrella mesoglea showing how these canals branch out over the entire bell. A dense carpet of white granules was found in the exumbrella mesoglea underneath radial canals. White scale bars in mm. Colored scale bar indicates relative OCT signal strength.

**Figure 6.**
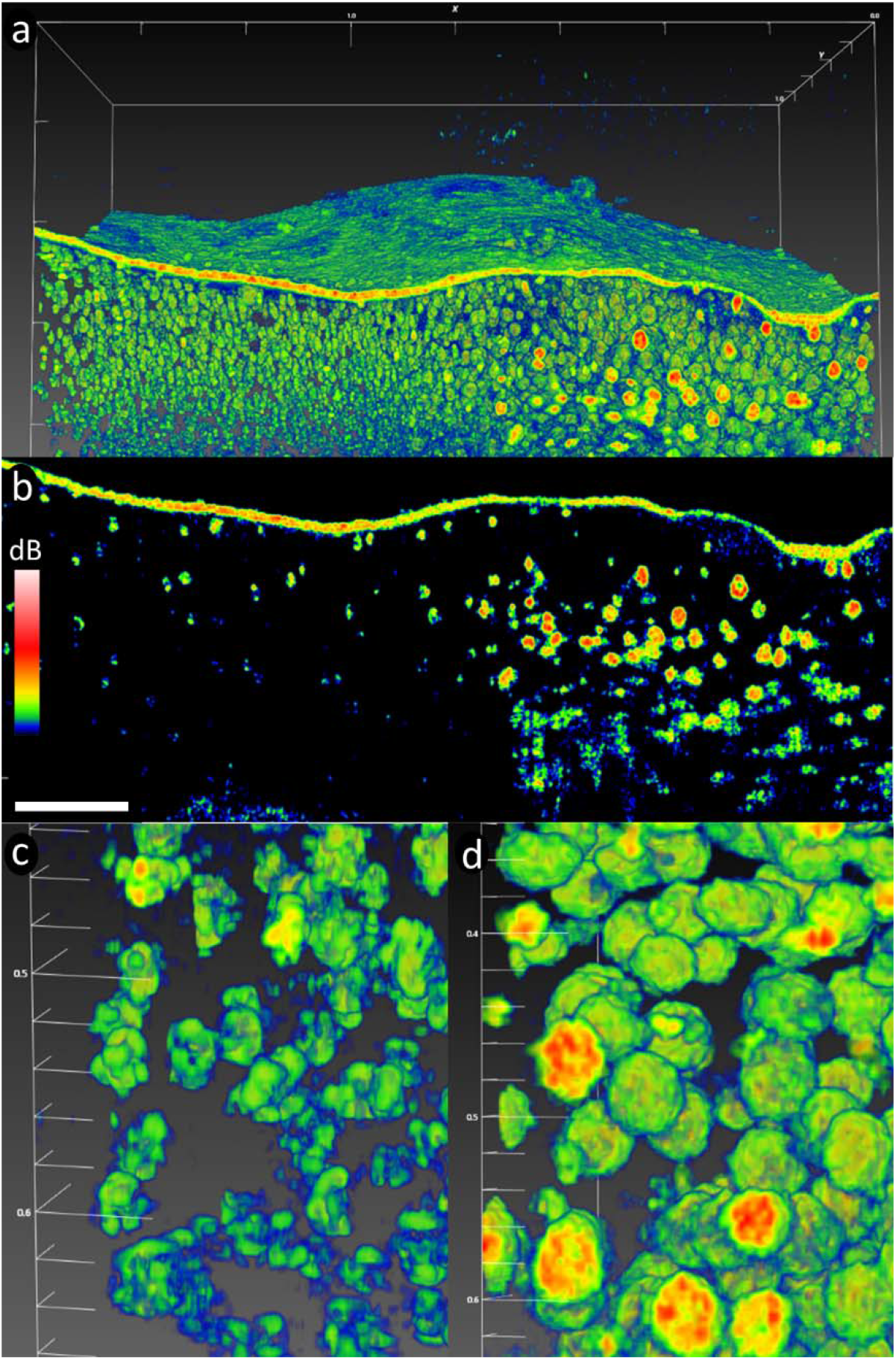
(a) 3D scan of a rhopalia canal with the subumbrella layer removed, revealing the density of symbiont clusters in anastomosing tissue (left side) *vs*. white granules (right side). (b) A sectional scan showing that white granules appear much denser than symbiont clusters. White granules were mainly spherical in shape (up to 80 μm in diameter), and appeared to be solid particles as judged from the OCT signal. (c) closeup-scan of symbiont clusters. (d) closeup-scan of white granules. Due to the density of granules, it was not possible to confidently discern symbiont clusters in this tissue region. White scale bars in panel a, c, and d in mm. White scale bar in panel b represents 200 μm. Colored scale bar indicates relative OCT signal strength. An animated 3D representation of panel a can be found in the supplementary information (Movie 2).

### Variable chlorophyll fluorescence imaging

The absorptivity and PSII quantum yield in the exposed bell tissue areas on each medusae were imaged with a variable chlorophyll fluorescence imaging system using blue LED’s (I-PAM, IMAG MIN/B, Walz; Ralph et al., 2005) providing weak, non-actinic measuring light pulses, strong saturating light pulses, as well as actinic light levels of known photon irradiance. All measurements of variable chlorophyll fluorescence were performed, and areas of interest (AOIs) drawn, by using the software ImagingWin (v2.41a, Walz).

Prior to imaging, medusae were individually dark acclimated for 15 minutes in a petri dish filled with ASW. During dark acclimation, the absorptivity of red light (a proxy for chlorophyll distribution) was measured as

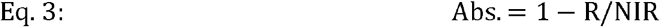

i.e., the fraction of red light absorbed by the sample relative to near infrared (NIR) light. The maximal PSII quantum yield (F_v_/F_m_) was then measured by applying a strong saturation pulse (>3000 μmol photons m^-2^ s^-1^ for 0.8 s). Rapid light curves (RLCs), i.e., relative PSII electron transport as a function of photon irradiance, were subsequently measured by measuring the effective PSII quantum yield, Y(II), and the relative electron transport rate, rETR (= Y(II) x PAR), at a range of increasing photon irradiance levels of photosynthetically active radiation, PAR (400-700 nm; 0-700 μmol photons m^-2^ s^-1^).

The surface areas of 6 AOIs on the exposed bell of each medusae were estimated with the built-in analysis function using the ImagingWin software. Pixels were converted to an estimated surface area:

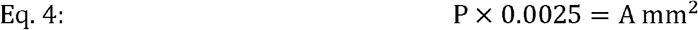

where P is the number of pixels in a given AOI, 0.0025 is the estimated conversion factor from the ImagingWin software, and A is the final area in mm^2^. The AOIs were chosen based on visible white granules along rhopalia canals and the anastomosing tissue connecting the canals.

### Isotopic pulse labelling experiment

#### Experimental setup

A tank with ∼10 L of deionized water was placed on top of 3 magnet stirrers (RCT basic, IKA GmbH). A heater and a small water pump were fitted in the bath to keep water homogenously at 25(±0.5)°C. A white LED lamp (KL2500 LED; Schott AG) equipped with a 3-furcated fiber guide was used for homogenous illumination over each magnet stirrer. Each fiber guide was fitted with a collimating lens, and light was adjusted to an incident photon irradiance of 350 μmol photons m^-2^ s^-1^ (400-700 nm), as measured with a calibrated spectroradiometer (MSC-15, GigaHertz Optik). The photon irradiance was chosen based on the average RLCs (Figure 9c), where all individuals showed light saturation above 500 μmol photons m^-2^ s^-1^. Isotopic labelling was done in 100 mL plastic beakers placed in the thermal bath and fitted with a magnet bar, as well as a mesh at the bottom to separate medusae from the rotating magnet (200 rpm).

#### Water preparation and pulse labelling

Prior to the labelling experiment, 1 L of isotopically enriched artificial seawater (ASW) was prepared as follows: Filtered (0.2 μm; Millipore) ASW (salinity of 35, pH 8.2) was stripped of dissolved inorganic carbon by lowering pH to < 3 with 1M HCl, and was then flushed with atmospheric air over night. The ASW was then enriched with 3mM NaH^13^CO_3_ (99 atom%, Sigma Aldrich), brought back up to experimental pH (8.2) with 1M NaOH, and 3μM ^15^NH_4_ Cl (99 atom%, Sigma Aldrich) was added. The mix was left to equilibrate with the experimental temperature of the thermal bath (25°C) before pulse labeling began.

Three *Cassiopea* medusae of ∼3.5-4 cm in diameter were chosen for the labelling experiment. For each medusa; 3-4 oral arms were removed using a tweezer and razor blade in order to expose one half of the medusa bell surface area. Medusae were subjected to 6 hours of pulse labeling in 80 mL of isotopically enriched ASW, with a change of isotopically enriched ASW every 2 hours. At the end of pulse labelling, one quarter of the bell from each medusae was chemically fixed in 10 mL of 2.5% [v/v] glutaraldehyde (Electron Microscopy Sciences), 4% [v/v] paraformaldehyde (Electron Microscopy Sciences), and 0.6 M sucrose (Sigma-Aldrich) mixed in 0.1 M Sörensen phosphate buffer. Chemically fixed *Cassiopea* samples were kept at room temperature for 2 hours and then stored at 4°C in fixative until further processing.

#### Histological sectioning for NanoSIMS analyses

A detailed description of sample preparations can be found in Lyndby et al. (2020). Briefly, isotopically labelled animals had a piece of the bell cut out including both anastomosing tissue and the rhopalia canal with visible white granules. The cut-out tissue samples were divided into the two respective tissue regions, and a smaller piece roughly halfway between manubrium and margin was extracted and embedded in Spurr’s resin. Semi-thin histological sections with a thickness of 200 nm were then cut with a Ultracut S microtome (Leica Microsystems), placed on 10 mm diameter round glass slips, and gold coated with a layer of 12.5 nm gold (Leica EM SCD050, Leica Camera AG) for NanoSIMS imaging.

#### NanoSIMS image acquisition

Isotopic imaging of semi-thin histological sections was performed with a NanoSIMS 50L instrument (Hoppe et al., 2013). Images (40×40 μm, 256×256 pixels, 5000 μs/pixel, 5 layers) were obtained with a 16 KeV Cs^+^ primary ion beam focused to a spot-size of about 120 nm (with a beam current of 2.2 pA). Secondary ions (^12^C_2_ ^-, 13^C^12^C^-, 12^C^14^N^-^, and ^12^C^15^N^-^) were counted in individual electron-multiplier detectors at a mass resolution of about 9000 (Cameca definition), sufficient to resolve all potential interferences in the mass spectrum (Clode et al., 2007; Gibbin et al., 2018; Lechene et al., 2006; Nuñez et al., 2018; Pernice et al., 2015).

Isotopic images were analyzed using the software L’IMAGE developed by Prof. Larry Nittler at the Arizona State University. Contours were drawn in each image around the epidermis as well as around individual amoebocytes and dinoflagellate cells. The epidermis in each NanoSIMS image was treated as a single region of interest (ROI; n = 1). Similarly, amoebocytes were treated as one ROI per image unless their cell clusters were clearly separated. Due to low numbers of symbionts in the exumbrella tissue, NanoSIMS analyses only include subumbrella tissue. Drift-corrected maps of ^13^C- and ^15^N-enrichment were obtained from the count ratios ^13^C^12^C^-^/^12^C_2_ ^-^ and ^15^N^12^C^-^/^14^N^12^C^-^, respectively. Measured enrichments were expressed in delta notation:

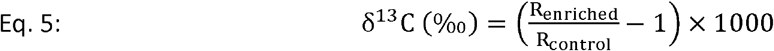

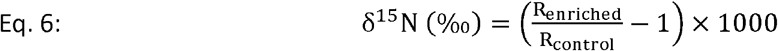

where R_enriched_ and R_control_ are the count ratios of an enriched sample and a control (i.e., unlabeled) sample, respectively. Total numbers of technical replicates (ROIs) are provided in Table S1.

### Symbiont density estimation

The density of symbionts was estimated from bell tissue corresponding roughly to AOIs drawn in ImagingWin (Walz GmbH). For each cut-out section, the subumbrella epidermis was carefully peeled off using a pair of fine tweezers, leaving the thick mesoglea (bulk mass of the bell) with the ex-umbrella epidermis still attached; simply referred to as “mesoglea” from this point. The subumbrella epidermis and the mesoglea were analyzed separately by homogenizing the tissue in 1.5 mL of filtered (0.45 μm) ASW using a T10 Standard Ultra Turrax handmixer (IKA) until samples were completely homogenized. Then 1 mL of the tissue slurry was transferred to a Sedgewick-rafter cell S52 (Pyser-SGI), and dinoflagellate cells were counted in 10 random squares on a microscope (AxioStar Plus, Zeiss). A total of 3 biological individuals were used, each of which 6 tissue regions were extracted (3 rhopalia canals and 3 anastomosing tissue regions).

### Statistical analyses

We used R (version 4.0.5) with the packages nlme (version 3.1-157) to perform statistical analyses. Linear mixed model (LMM) analyses were used to test the relationship between isotopic enrichment in *Cassiopea* holobiont compartments in 2 bell tissue regions (near rhopalia canals and in anastomosing tissue) taking into account the biological replicate as a random factor. ^13^C enrichment data for amoebocytes, and ^15^N enrichment data for symbionts and host epidermis were square root transformed to achieve normality.

## Results and Discussion

### Imaging *Cassiopea* morphology

Due to the near-neutral buoyancy of *Cassiopea* jellyfish, even small vibrations during OCT imaging can cause the sample to move, creating distortion in the scans. Scanning speed and reference light intensity was thus adjusted to achieve a relative short scan-duration without sacrificing signal intensity. The field of view (FOV 7×7×2.8 mm; X-Y-Z) was adjusted to contain the entire animal, and the number of pixels in each direction was kept to a minimum (350×350×1024 pixel; X-Y-Z) (Figure 1a). The scan pattern resulted in a voxel size of 20×20×2.74 μm (X-Y-Z). Given that the average diameter of dinoflagellate symbionts found in *Cassiopea* are typically around 10 μm (Biquand et al., 2017; Fitt, 1985), this voxel size provided sufficient resolution to identify symbiont clusters in form of amoebocytes (that typically host >10 cells per unit, e.g., Lyndby et al., 2020), even from complete specimen scans (Figure 1b).

Externally, rhopalia were visible along the bell margin, and the first oral arm bifurcation and groves were clearly visible. Individual arms showed signs of vesicles forming apically, while digitata were not yet present (e.g., Jordano et al., 2022; Muffett et al., 2022) (Figure 1a). The bell-disk surface is not perfectly flat, but appears ridged in a radial pattern, possibly due to the musculature lining the subumbrella epidermis (Blanquet and Riordan, 1981). Some shading is apparent from oral arms and the marginal lobe reaching in over the bell disc, and the OCT generally did not manage to penetrate through the entire animal system from sub-to exumbrella epidermis due to multiple scattering effects degrading the contrast with depth. Internally, symbiont clusters and white granules were found in high densities near the subumbrella epidermis (e.g., Estes et al., 2003; Lyndby et al., 2020), and white granules can be seen extending further down into the exumbrella mesoglea along rhopalia canals (Figure 1b).

Changing to a higher magnification objective, it was possible to zoom in on the above-mentioned structures to reveal more details, while also gaining more non-invasive information on the internal tissue organization of *Cassiopea*. We used both intact animals and cut-out tissue regions from multiple specimens of various sizes to scan a broad range of anatomical structures at different growth stages.

Rhopalia are specialized eyespots in Medusozoa that allow medusa to detect light and orientate themselves in the water column (Kayal et al., 2018; Martin, 2002). *Cassiopea* have on average 16 rhopalia distributed along the bell margin (Bigelow, 1900). Closeup scans with OCT showed that these eyespots are well nested between two marginal lappets along the rim (Figure 2a-c). Rhopalia canals are major canals of the gastric network extending from the center of the bell, out towards the rhopalia (Bigelow, 1900) (Figure 2d, e). Cross sectional scans near one of the rhopalia showed that these major canals fork into two canals, each reaching out into a marginal lappet around the rhopalium (Figure 2d, e).

Oral arms are highly branched at the center of the bell (Figure 3a). They typically occur in four pairs in *Cassiopea*, and each arm harbors secondary mouths and vesicular appendages. Vesicular appendages could be seen containing clusters of cassiosomes ready for deployment to catch prey (Ames et al., 2020) (Figure 3b). Cassiosomes are snared off from the vesicular epidermis in amorphous ‘popcorn’ shaped tissue balls, with newly developing cassiosomes being formed underneath fully developed ones (Ames et al., 2020). However, it was not possible to confidently discern individual cassiosomes underneath the surface of the cluster, possibly due to the accumulated layers of dermis from each individual cassiosome that substantially lowered OCT scan penetration depth into these clusters (Figure 3b).

### Improving depth penetration

During this OCT imaging work, several factors that decreased the signal penetration were identified. Host epidermal tissue, symbionts, and white granules all greatly scattered the NIR light from the OCT light source decreasing the contrast with depth in the jellyfish tissue. To reduce the OCT signal loss at the upper tissue interface, the refractive index of the surrounding medium was matched to the epidermis in *Cassiopea*. Glycerol has a refractive index of 1.473, and submerging a piece of tissue in pure glycerol demonstrated that the *Cassiopea* epidermis has a very similar refractive index, thus permitting OCT imaging of deeper tissue layers (Figure 4). Sectional scans of a vesicular lappet clearly showed the entire structure, and revealed a cavity in the center of the lappet (Figure 4b). Contrary to tissue scanned in water, where the endodermis/gastrodermis appeared thin or transparent, the endodermis of the lappet cavity appeared much clearer due to less backscattering in the epidermis. Individual symbiont cluster also became more distinctly visible throughout the lappet system when imaged in pure glycerol (Figure 4).

While pure glycerol matches the refractive index of the epidermis well, the high viscosity of glycerol makes it difficult to image larger samples, and submerging intact medusae in glycerol did not improve depth penetration due to water and mucus sticking to the surface of the animal, which resulted in lower signal penetration and contrast.

To explore the deeper layers of the bell, we cut a piece of bell tissue from a medusa and then carefully removed the subumbrella epidermis and mesoglea using a pair of fine tweezers. Removing the subumbrella epidermis greatly improved the OCT depth penetration, because the symbiont cell clusters and the epidermis were the main sources of light scattering in these anatomical structures (Figure 5a). When the tissue was cut, grabbing and ripping off the subumbrella epidermis proved easy, and even kept parts of the exumbrella gastrodermis from radial canals still visibly attached to the exumbrella mesoglea (Figure 5b).

Removing the subumbrella epidermis greatly improved visibility of symbiont clusters and white granules present in the exumbrella mesoglea. A side-by-side comparison of the exumbrella mesoglea, with half the scan covering the rhopalia canal full of white granules, and the other half covering anastomosing tissue only with symbiont clusters, revealed a major heterogeneity in the structure and content of the two tissue areas (Figure 6).

### High-resolution scans

High-resolution scans with a voxel size of 0.5×0.5 (Figure 6c) or 1.0×1.0 μm area (Figure 6d), and 2.7 μm in height, were used for detailed scans of the spatial distribution of symbiont clusters and white granules. White granules were found in high density along the rhopalia canal, and appearing to be ‘solid’ spheres of roughly 80 μm in diameter (Figure 6b). Symbiont clusters were also found in surprisingly high density in the exumbrella epidermis, although more diffusely spread out compared to the white granules. Symbionts appeared to be positioned in elongated clusters, stretching up to 80 μm across. Given that the average size of *Symbiodinium* cells *in hospite* with *Cassiopea* are normally around 10 μm in diameter (Biquand et al., 2017; Fitt, 1985) this indicates that amoebocytes can harbor more than 10 symbionts depending on clustering formation, consistent with a previous estimate (Lyndby et al., 2020). Due to the high density of white granules near rhopalia canals, clusters of symbionts were hard to detect or distinguish from the granules in this region (Figure 6d). However, symbiont densities quantified from the two regions by cell counting were found to be equally distributed between the anastomosing- and rhopalia canal tissue (Figure 7a). We also found that the symbiont populations were more than 3-fold higher in the subumbrella layer relative to the exumbrella layer (Figure 7b). The data are in agreement with the larger OCT scans of intact tissue, which showed that symbionts were more abundant near the subumbrella epidermis, as compared to the rest of the *Cassiopea* medusa (Figure 1b).

**Figure 7.**
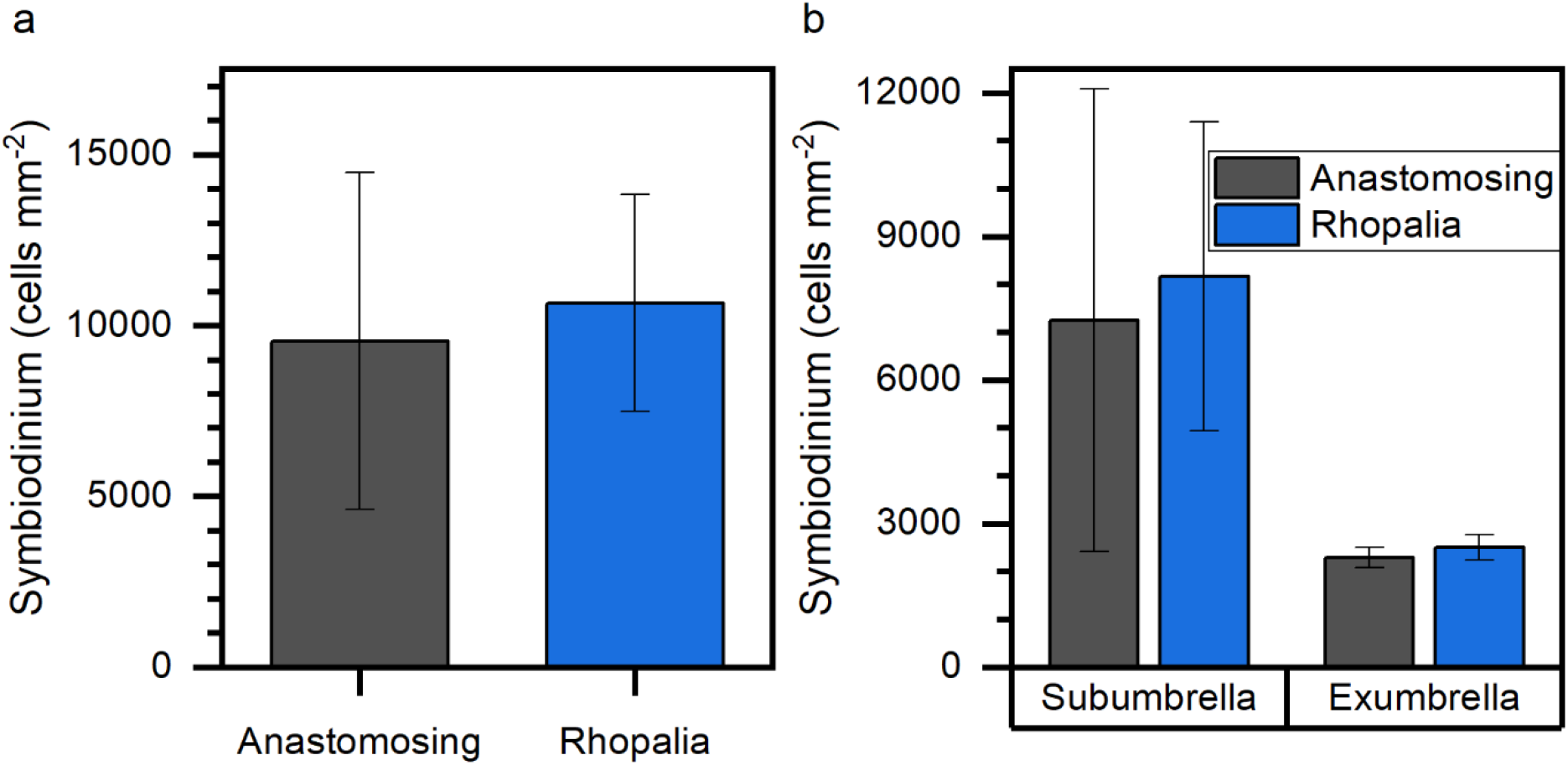
(a) Symbiont densities in anastomosing tissue and rhopalia canals averaged over the entire depth of the bell. (b) Symbiont densities observed within the anastomosing tissue and rhopalia canals but distinguishing between the subumbrella and exumbrella layer of the endodermis. Densities were averaged against the estimated surface area of tissue cut-outs (see methods). Data show means ±SE (n = 3 biological replicates).

### Optical properties of *Cassiopea* bell tissue

Studies of the optical properties of cnidarian tissue is an expanding field, yielding important information about how host tissue and symbionts modulate and interact with their light environment. Such information is also important for the development of models that simulate radiative transfer e.g. in corals (e.g., Taylor Parkins et al., 2021). While the function of the white granules in *Cassiopea* remains unknown, other pigments in *Cassiopea* have been speculated to be either photoprotective or photo-enhancing (Blanquet and Phelan, 1987; Phelan et al., 2006), similar to host pigments found in corals (e.g., Lyndby et al., 2016; Salih et al., 2000). The high density of the heterogeneously distributed light scattering white granules in the bell and oral arms suggests that the light microenvironment could be greatly altered in these tissue regions relative to tissue with no white granules. We estimated the optical properties of *Cassiopea* bell tissue with light scattering white granules using optical parameter extraction from OCT scans (Wangpraseurt et al., 2019).

The optical properties of tissue with white granules near the rhopalia canals were radically different from the anastomosing tissue (Figure 8). The scattering coefficient of tissue with white granules were estimated to be μ_s_ = 200-300 cm^-1^, and with an anisotropy factor of *g* = 0.7 for individual white granules (i.e., analyses targeting narrow regions with individual granules; Fig. S1a). Estimates from larger areas of tissue (i.e., analyses averaged over large areas, comprising multiple granules with a width of 10-20 granules; Fig. S1b) indicated that *g* increased to 0.8-0.9 and the scattering coefficient dropped to μ_s_ = 40-100 cm^-1^. This suggests that the arrangement of light scattering white granules in the tissue increases both the photon residence time and depth penetration, thereby enhancing the chance of absorption by symbionts near white granules, similar to reflective (and fluorescent) host pigments in corals (Lyndby et al., 2016; Wangpraseurt et al., 2019).

**Figure 8.**
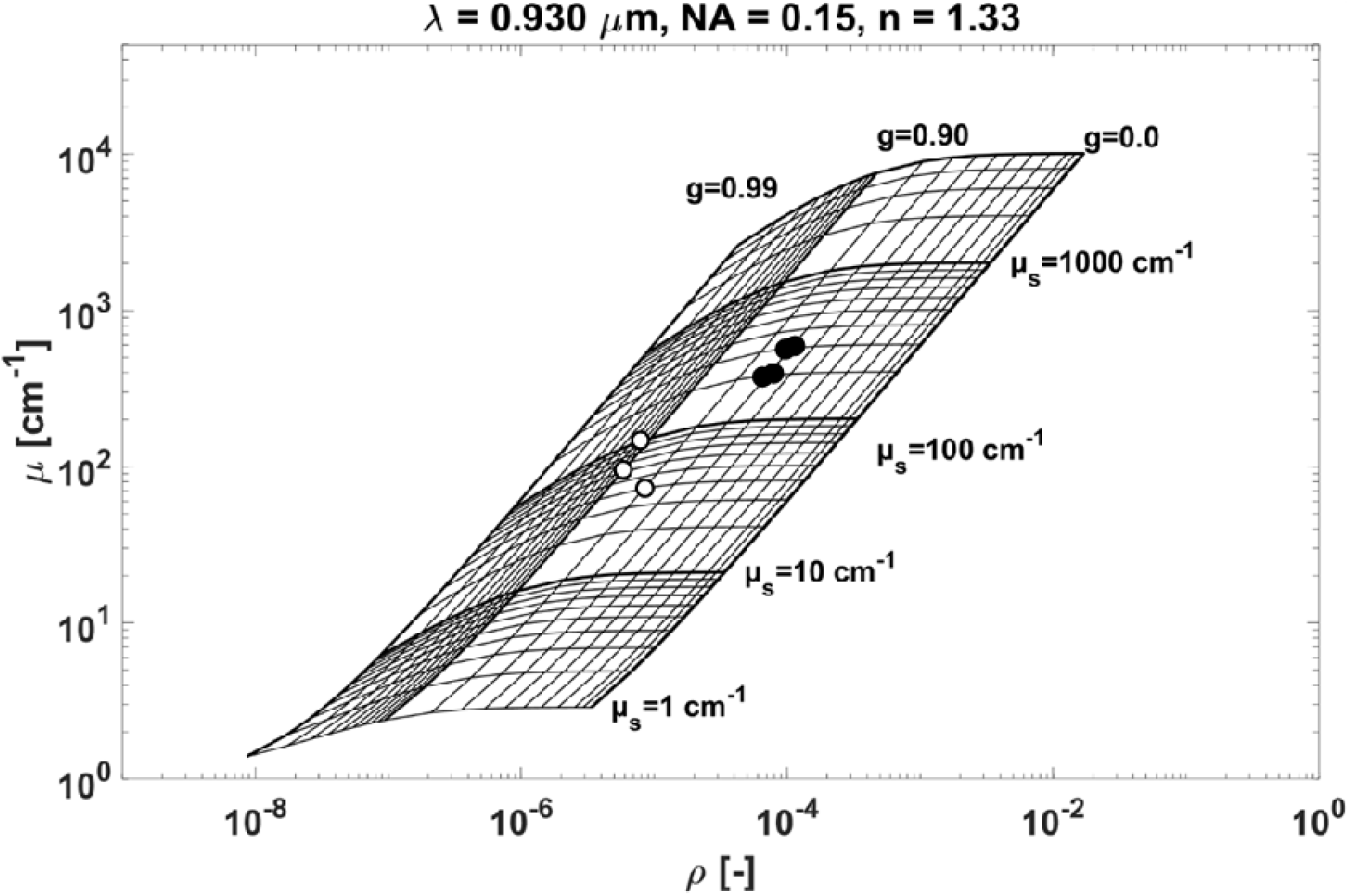
The optical properties, i.e., scattering coefficient (μ_s_), and anisotropy of scattering (*g*), extracted from OCT scans of narrow regions targeting individual white granules (full circles) or large areas comprising of multiple granules (width of 10-20 granules per region; open circles) in rhopalia canal tissue with epidermis removed.

**Figure 9.**
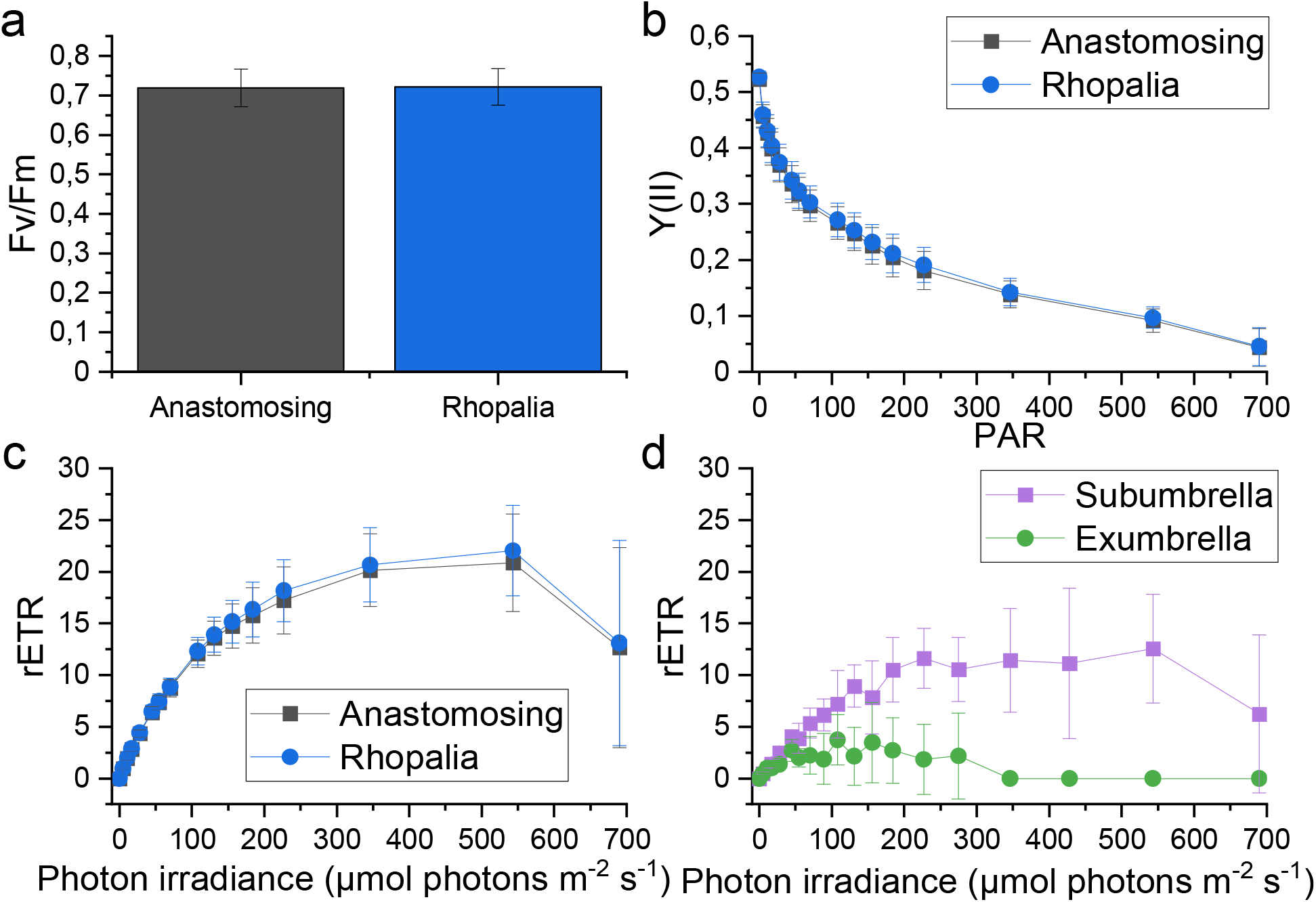
Variable chlorophyll fluorescence imaging of anastomosing tissue and rhopalia canals. (a) Maximum quantum yield (F_v_/F_m_) after 15 min dark adaptation, (b) effective PSII quantum yield [Y(II)], (c) relative electron transport rate (rETR) of intact medusae, and (d) rETR measured on cut-out bell tissue (from different medusa) with subumbrella and exumbrella layers separated as a function of incident photon irradiance. Data show means ± s.e.m. (n = 3 biological replicates for panel a-c, and n = 1 biological replicate for panel d).

It was not possible to determine the optical properties of symbiont clusters in anastomosing tissue. Symbiont densities in the exumbrella mesoglea were on average 3-fold lower than in the subumbrella, with few clusters that were diffusely spread in the mesoglea, as judged from B-scans of this tissue region (Figure 6b, c). This prevented the extraction of meaningful optical properties, but indicates that light travels through the exumbrella mesoglea in anastomosing tissue relatively unhindered. These differences suggest that symbionts in mesoglea with white granules might be more light-exposed based on an increased light scattering in this region, as compared to other tissue regions.

### Photosynthetic performance in different *Cassiopea* tissue compartments

Given that the optical properties of light scattering white granules can drastically alter the local light microenvironment, a 6-hour isotopic labelling experiment was conducted in combination with NanoSIMS isotopic analyses (Kopp et al., 2015; Krueger et al., 2017; Lyndby et al., 2020; Pernice et al., 2012). We used ^13^C-bicarbonate and ^15^N-ammonium incubations to investigate whether carbon and nitrogen assimilation rates differed between symbiont algae in the subumbrella anastomosing tissue and rhopalia canal regions. We combined such measurements with variable chlorophyll fluorescence imaging to investigate the photosynthetic performance and carbon assimilation of symbionts in the two tissue regions.

PAM chlorophyll fluorescence imaging showed no differences in F_v_/F_m_ between algal symbionts found along rhopalia canals *vs*. anastomosing tissue in intact bell tissue (Figure 9a), and the effective quantum yield was also similar for the two regions, as measured with RLCs (Figure 9b). However, removing the subumbrella layer, and running RLCs on the sub- and exumbrella layers side-by-side showed that symbionts deeper in the bell had a drastically lower threshold for light saturation, and rETR saturated already around 100-200 μmol photons m^-2^ s^-1^ (Figure 9d), relative to saturation at 400-500 μmol photons m^-2^ s^-1^ for symbiont photosynthesis in the subumbrella layer (Figure 9c, d). This indicates that symbionts in the exumbrella epidermis are acclimated to lower light levels, suggesting that most light is absorbed at the subumbrella epidermis directly, or is diffused/scattered by the white granules.

Stable isotope labelling of three sub-adult specimens of *Cassiopea* showed a significant difference in the ^13^C assimilation by symbiont algae near white granules, relative to symbionts found in anastomosing tissue (LMM, F_1,4_ = 8.8, *p* = 0.041; Figure 10a), suggesting that photosynthesis is locally enhanced by the light scattering from white granules. No differences in ^13^C enrichment were observed in amoebocyte cells (LMM, F_1,4_ = 1.2, *p* = 0.342) or host epidermis (LMM, F_1,4_ = 0.5, *p* = 0.508). Finally, no differences in ^15^N uptake were found for symbionts (LMM, F_1,4_ = 0.04, *p* = 0.854), amoebocytes (LMM, F_1,4_ = 0.5, *p* = 0.508), or host epidermis (LMM, F_1,4_ = 0.2, *p* = 0.649), comparing between rhopalia canals and anastomosing tissue (Figure 10b), indicating that each region had equal access to nutrients during the labelling experiment.

**Figure 10.**
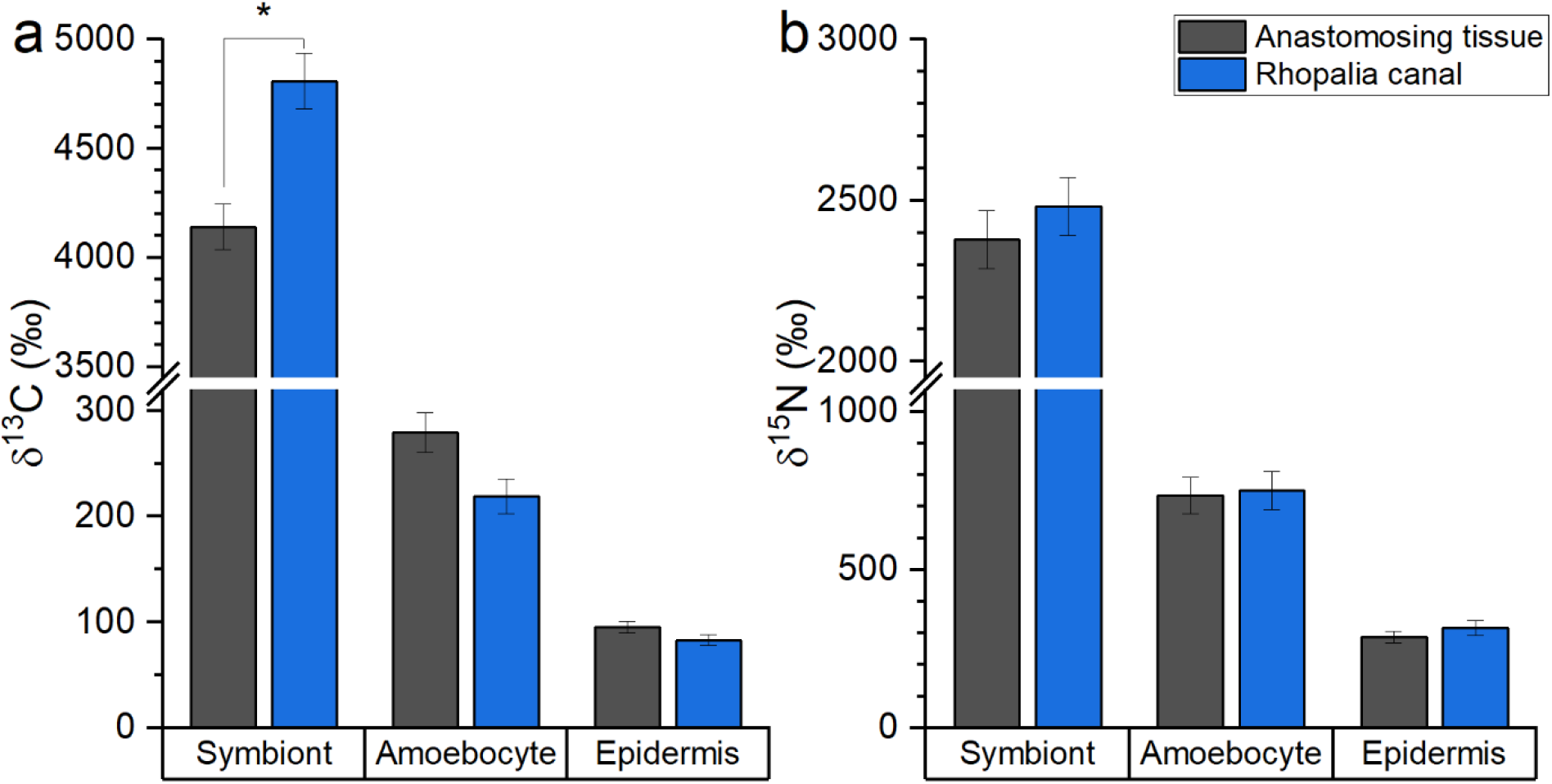
Mean ^13^C- (a) and ^15^N enrichments (b) of symbiont algae, amoebocyte cells, and host epidermis in subumbrella tissue in 3 sub-adult *Cassiopea* medusae after a 6-hour pulse labelling incubation with ^13^C-bicarbonate and ^15^N-ammonium. * indicates significant difference at the level p < 0.05 between respective tissue/cell types from the two tissue regions. Error bars indicate ± s.e.m. (n = 3 biological replicates).

The presence and role of optical microniches in *Cassiopea* for symbiont photosynthesis has hitherto been unexplored. Here, we provide evidence that the *Cassiopea* host directly alters the light microenvironment, benefitting nearby symbionts. White light scattering granules in *Cassiopea* provide a means for the host to not only enhance photosynthetic performance of symbionts, but also to propagate light further down into the host tissue, potentially supporting symbiont populations present in shaded parts of the animal. While it remains unknown what triggers production of white light scattering granules in *Cassiopea*, it is possible that individuals living in overall more shaded environments, e.g., mangroves, would be more inclined to produce white light scattering granules to enhance photosynthesis of symbionts.

We found highest symbiont densities near the subumbrella epidermis due to higher light availability. However, enhanced propagation and homogenization of light through the *Cassiopea* tissue via scattering might explain why roughly one quarter of the symbiont population is found deeper into the bell and exumbrella layers. Furthermore, light penetrating all the way through the animal to the substrate could potentially be backscattered up into the animal, additionally enhancing photon availability in the umbrella.

## Conclusion

Optical coherence tomography can provide detailed scans of the *in vivo* tissue organization of intact *Cassiopea*, showing the distribution of symbiont clusters, white granules, and other structures at high spatial resolution. The methods presented here enable detailed mapping of the distribution of entire symbiont populations or similar distinguishable objects in intact, living cnidarian systems, providing new means for non-invasive monitoring of the internal dynamics of symbiotic cnidarians, under both homeostasis and during stressful events. Additionally, this study provided the first insight into how light scattering granules in *Cassiopea* medusae can enhance the photosynthetic performance of their symbionts near these granules. These observations support previous suggestions that white tissue regions with granules in *Cassiopea* play several roles in modulating the light microenvironment of *Cassiopea*, by enhancing light availability in through scattering, thus exhibiting both photo-enhancing and photoprotective properties. The presented approach to study structure and function in *Cassiopea* is also applicable to other photosymbiotic cnidarians (e.g., cnidarian model systems such as *Exaiptasia* sp. and *Hydra* sp.), but we propose that OCT could also find broad application for non-invasive monitoring of structure and morphology of particular tissues or whole specimens in many types of invertebrates.

## Acknowledgements

The authors thank the staff at the Electron Microscopy Facility at University of Lausanne for excellent support and assistance with processing samples for NanoSIMS analyses.

## Competing interests

The authors declare no competing interests.

## Author contribution

NHL, SM, AM, and MK designed the experiment. NHL, SM, SB, and SLJ carried out experiments and data collection. NHL, SM, MK, and AM analyzed the data. All authors contributed to writing of the manuscript.

## Funding

This study was supported by an award from the Gordon and Betty Moore Foundation (grant no. GBMF9206; https://doi.org/10.37807/GBMF9206 to M.K), and the Swiss National Science Foundation (grant no. 200021_179092 to A.M.).

## Supplementary information

**Table S1.** NanoSIMS isotopic enrichment data summary. Treatment IDs correspond to the 2 bell regions detailed in the methods section of the main text. Delta values of ^13^C and ^15^N represent mean values ± s.e.m. calculated from all replicates of a given treatment/tissue area as described by equation 4 and 5 in the methods section.

| Bell region | Tissue/cell type | Replicates | $^{13}\text{C}$ | $^{15}\text{N}$ |
| --- | --- | --- | --- | --- |
| Anastomosing | Symbionts | 218 | $3798.3 \pm 122.9$ | $2181.8 \pm 94.0$ |
| Anastomosing | Amoebocytes | 45 | $236.2 \pm 22.0$ | $621.4 \pm 62.4$ |
| Anastomosing | Epidermis | 39 | $74.1 \pm 7.5$ | $222.3 \pm 24.1$ |
| Rhopalia canal | Symbionts | 203 | $4192.9 \pm 158.8$ | $2163.3 \pm 97.4$ |
| Rhopalia canal | Amoebocytes | 45 | $180.1 \pm 18.5$ | $617.7 \pm 65.8$ |
| Rhopalia canal | Epidermis | 37 | $64.4 \pm 7.2$ | $246.8 \pm 28.6$ |

**Fig. S1.**
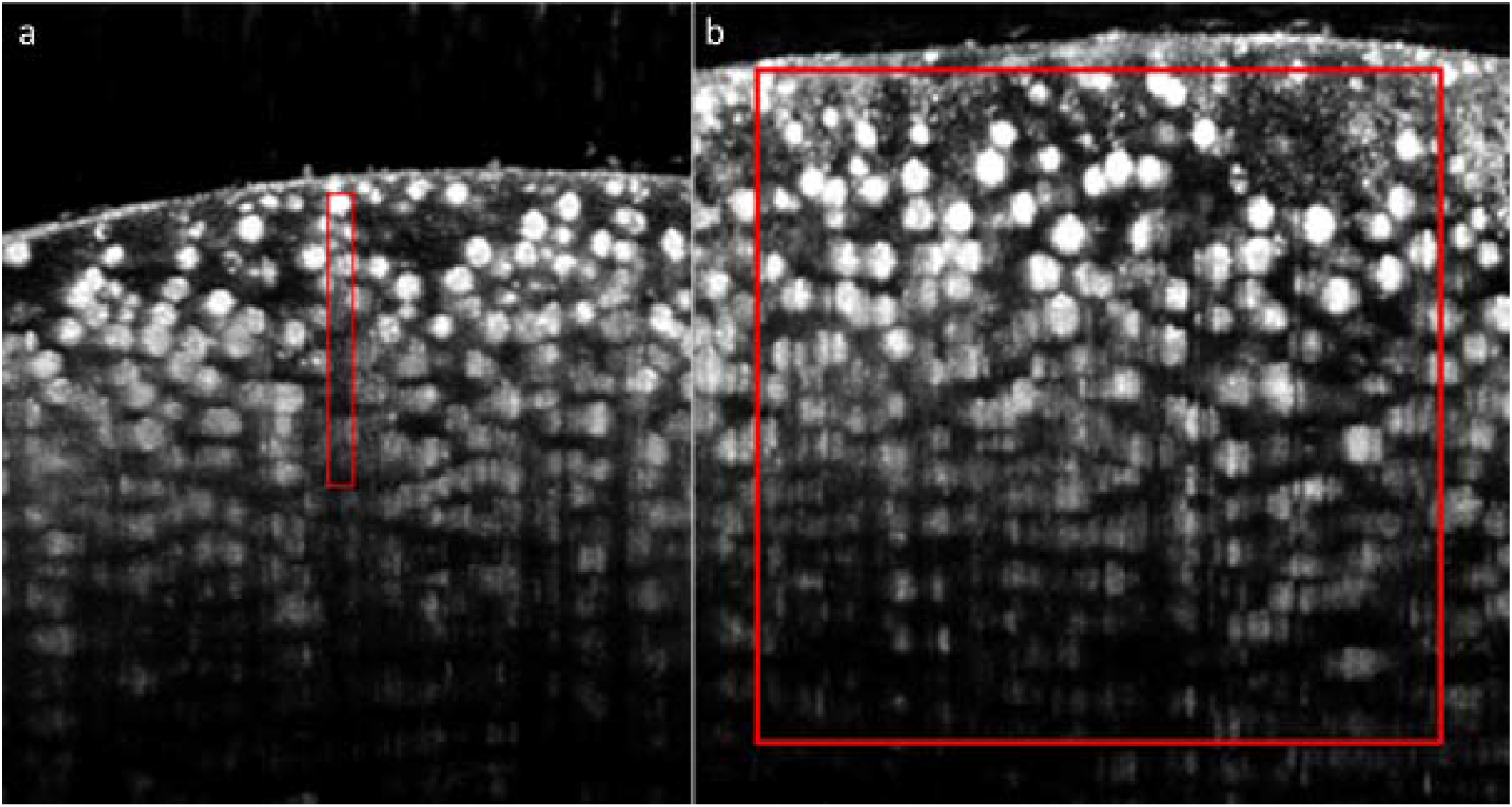
Example of rhopalia canal tissue regions for inherent optical parameter extraction, showing the region for (a) individual white granules, and (b) large area with multiple granules (width of 10-20 granules).

Movie 1 | Animation of 3D scan of an entire juvenile *Cassiopea* sp. medusa (6 mm diameter) using a FOV of 7×7×2.8 mm (X-Y-Z), a resolution of 350×350×1024 pixels, giving a voxel size of 20×20×2.74 μm.

Movie 2 | Animation of 3D scan of exumbrella tissue after removal of the subumbrella layer to increase depth penetration and clarity. Scan was done right at the transition between rhopalia canal tissue, with a high presence of white light scattering granules (larger spheres), and the anastomosing tissue with a dense population of symbionts clustered in amoebocytes (smaller, elongated spheres) and no visible white light scattering granules, showing a clear division of the two types of tissue.

## Notes

### Competing Interest Statement

The authors have declared no competing interest.

